# Dynamic cholinergic signaling differentially desynchronizes cortical microcircuits dependent on modulation rate and network topology

**DOI:** 10.1101/2025.06.20.660675

**Authors:** Sibi Pandian, Scott Rich

## Abstract

Acetylcholine (ACh) affects both the intrinsic properties of individual neurons and the oscillatory tendencies of neuronal microcircuits by modulating the muscarinic-receptor gated m-current. However, despite contemporary experimental evidence of ACh concentrations changing at millisecond timescales, computational studies traditionally model ACh solely as a tonic neuromodulator. How time-varying, dynamic cholinergic modulation of the m-current affects the dynamics of neuronal microcircuits therefore remains an open question. Using a new implementation of a time-varying cholinergic signal in computational excitatory-inhibitory (E-I) spiking neuronal networks, we here delineate how the interaction between dynamic cholinergic modulation and network topology influences the oscillatory tendencies of these systems. While the dynamics of networks with dominant inter-connectivity (strong E-to-I and I-to-E synaptic weights) are minimally affected, networks with dominant intra-connectivity (strong E-to-E and I-to-I synaptic weights) exhibit dynamics heavily dependent upon dynamic cholinergic signaling. Further investigation of these latter type of networks reveals that their firing patterns are sensitive to the timecourse of cholinergic modulation and that relatively minor changes to the E-I connectivity strength promote distinct desynchronization mechanisms. Our results indicate that network topology plays a paramount role in dictating the modulatory effects of time-varying cholinergic signals, a finding of broad relevance to our understanding of cholinergic modulation and potentially impactful in the design of neurostimulation therapies believed to act through cholinergic pathways.

**Author summary:** Acetylcholine (ACh) is a chemical messenger that alters the intrinsic properties of neurons and in turn regulates brain activity related to cognitive functions such as sensory processing, memory formation, and attention. Although mounting experimental evidence shows that ACh concentrations in the brain can change dynamically at rapid timescales, computational studies conventionally consider ACh concentrations as constant across time. This motivated us to create a computational model of a neuronal microcircuit in which cellular properties are affected by ACh concentrations varying at millisecond-level timescales. The oscillatory dynamics of this microcircuit differ in important ways from similar systems with constant ACh levels, with the influence of dynamically changing ACh critically dependent upon the connectivity of the neurons in the modeled brain region. These results describe novel mechanisms by which ACh controls microcircuit activity overlooked in existing models in which ACh varies only over long timescales, an understanding which may be vital for the refinement of neurostimulation therapies believed to act through transient alterations to ACh activity.

## Introduction

Acetylcholine’s (ACh’s) important role in the regulation of brain activity has traditionally been contextualized via its influence as a tonic neuromodulator acting over long timescales. ACh modulates individual neurons’ cellular excitability by binding to muscarinic receptors, correspondingly exerting an influence on the firing patterns of neuronal microcircuits [1]. Functionally, ACh plays a role in the facilitation of cognitive processes such as attention, sensory processing, and memory formation [2]. High levels of ACh are present during wakefulness and low levels during sleep, a phenomenon that may contribute to the desynchronization or decorrelation of cortical activity during awakening as observed in electroencephalogram recordings [3, 4]. Its significance for cognitive function is also reflected in pathology: deterioration of the basal forebrain, the primary source of cholinergic projections to the cortex, correlates with the onset of Alzheimer’s disease and other instances of cognitive decline [5].

Although cholinergic signaling to the cortex is conventionally viewed as slow and diffusive—acting over near-tonic timescales at least as long as seconds—recent technological advancements have enabled measurement of ACh concentrations with higher temporal resolution and challenged this notion [6, 7]. *In vivo* studies have demonstrated the presence of cholinergic transients in the cortex, initiating and peaking at timescales of hundreds of milliseconds or shorter, in response to relevant stimuli [8]. These cholinergic transients play a key role in perceptual tasks that demand immediate reactions [1, 2, 5], such as cue-detection [9] and foot shock-reactions [10]. Furthermore, recent evidence has emphasized that muscarinic pathways may play a critical role in facilitating rapid cholinergic control mechanisms of cortical circuits, especially in deeper cortical layers [11].

Computational modeling has proven effective in delineating the effects of cholinergic modulation at the cellular level and relating them to mesoscale neuronal microcircuit dynamics. In conductance-based neuron models, the neuromodulatory effects of ACh can be represented through closure of the m-channel [12] and thereby the m-current, an amalgam of multiple muscarinic-receptor gated slow potassium currents [13, 14] reported in experimental literature. Blockade of this current causes changes in cellular properties expected to arise from cholinergic modulation, including increased excitability, altered phase response curve (PRC) characteristics, and decreased spike frequency adaptation [12, 15–17].

Computational studies of cholinergic modulation on microcircuit dynamics have ranged in focus from populations of inhibitory interneurons [18] to populations of excitatory pyramidal cells with nonuniform ACh distribution [19]. Most common, though, are studies focusing on cholinergic modulation in excitatory-inhibitory (E-I) networks, networks consisting of inter-connected excitatory and inhibitory cell populations. E-I networks are of particular interest as they have been used in seminal computational work to model generic cortical microcircuits [20, 21], and have led to the proposal of the Pyramidal Interneuron Network Gamma (PING) mechanism [22]—an important conceptual model for the production of oscillatory network dynamics [23] based on the mutual entrainment of excitatory and inhibitory populations. Recent studies in E-I networks have characterized functional properties of cholinergic modulation [24–28] and also identified that features of network topology strongly influence the response to ACh [29]. However, with very few exceptions [28], these studies predominantly consider tonic cholinergic modulation via changes to a static m-channel conductance.

To address this gap, we investigated how a dynamic model of cholinergic modulation of the m-current affects the oscillatory properties of E-I networks, aiming to identify and explain unique effects of phasic cholinergic signaling. We compared the effects of tonic modulation with dynamic modulation across differing connectivity schemes and varying rates of modulation. Our explorations reveal that dynamic cholinergic modulation preferentially affects networks where intra-connectivity (E-to-E and I-to-I synapses) dominates inter-connectivity (E-to-I and I-to-E synapses). We probed these models to propose specific mechanisms by which dynamic cholinergic modulation disrupts network oscillations, finding that said mechanisms are surprisingly sensitive to minor alterations in the strength of the E-I synapses. These *in silico* results provide precise insights into how network topology mediates the effects of millisecond-scale cholinergic signals on network dynamics.

## Materials and methods

### Neuron Model

We model all excitatory and inhibitory cells in our E-I network with a Hodgkin-Huxley formalism originally used for cortical pyramidal neurons in [12, 17]. Although these equations were originally developed to model excitatory pyramidal cells, they have also been used to model inhibitory interneurons [18].

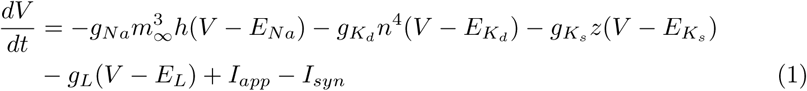

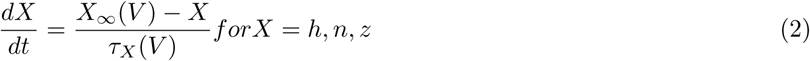

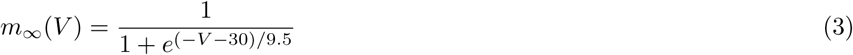

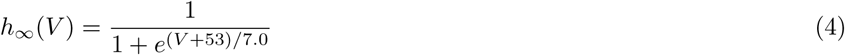

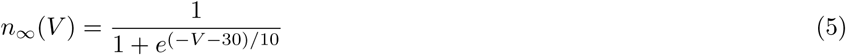

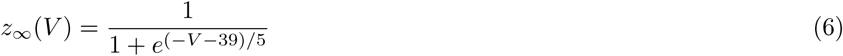

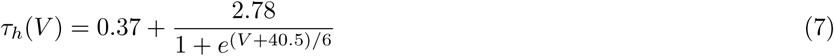

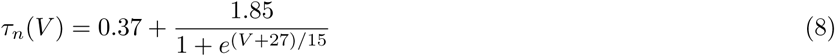

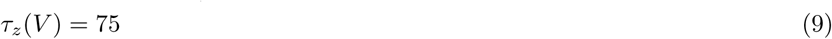

*V* represents the voltage of the neuron in mV, while *m*, *h*, *n*, and *z* represent the unitless gating variables of the ionic current conductances. *I_app_* and *I_syn_* signify the external applied current and the synaptic current (defined in Network Structure) respectively in *µ*A*/*cm^2^. 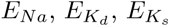, and *E_L_* are the reversal potentials, and 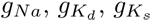, and *g_L_* the maximum conductances of the sodium channel, delayed rectifier potassium channel, slow m-type potassium channel, and leak-channel respectively. We use 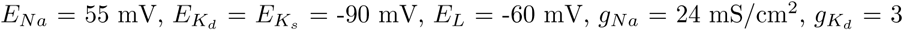 mS/cm^2^, and *g_L_* = 0.02 mS/cm^2^.

We model dynamic cholinergic modulation’s effects on cellular properties by subjecting 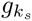 to a linear decline from 1.5 to 0 mS/cm^2^ for all cells, after allowing dynamics to initially stabilize at the maximal 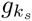 for 1000 ms.

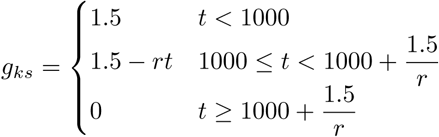

Here, *t* refers to the time in ms and *r* refers to the rate of linear decline in mS/cm^2^/(ms). We also study scenarios where we permanently set the inhibitory cells’ 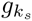 to 0 mS/cm^2^, only carrying out modulation with the excitatory cells.

### Network Structure

Our E-I network consists of 1000 neurons, 800 excitatory and 200 inhibitory. We establish inter-connectivity by allowing each excitatory cell a 50% chance to synapse onto each inhibitory cell, and each inhibitory cell likewise a 50% chance to synapse onto each excitatory cell. Additionally, we establish intra-connectivity by allowing each cell in the excitatory and inhibitory populations to have a 30% percent chance of synapsing onto other cells within their own population. We modeled synaptic connections using a counductance-based double-exponential profile of the form

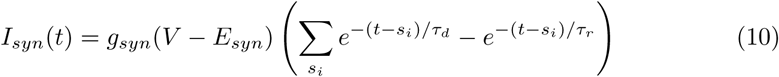

where *g_syn_* is the maximum synaptic conductance, *V* is the membrane voltage of the post-synaptic neuron, *E_syn_* is the reversal potential of the synaptic current, *s_i_* are the times of all presynaptic spikes occurring before the current time t in ms, and *τ_d_* and *τ_r_* are the synaptic decay and synaptic rise constants respectively. We set *E_syn_* to −75 mV for inhibitory synapses and to 0 mV for excitatory synapses. *τ_r_* is set to 0.2 ms for all synapses, while *τ_d_* is 3.0 ms for excitatory synapses and 5.5 ms for inhibitory synapses. To vary network topology, we control the strengths of the E-E, E-I, I-E, I-I synapses by individually altering the synaptic weights (*g_syn_*) for each type of synapse.

To introduce cellular heterogeneity, we choose *I_app_* for each excitatory cell such that, in isolation, the cell would fire at a randomly selected frequency within the uniform distribution (45 Hz, 55 Hz). Even as 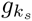 declines and cellular properties are changed, we concurrently modulate *I_app_* for every 0.01 mS/cm^2^ change in 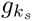 to preserve the originally selected firing frequency of the excitatory cells. Meanwhile, the inhibitory cells’ firing is driven largely by the excitatory signaling. *I_app_* for the inhibitory population is set such that the cells are brought closer to their threshold for firing, but does not by itself permit them to spike. Here too, we incorporate cellular heterogeneity and select *I_app_* for each cell from a (0.95 *I_A_*, 1.05 *I_A_*) uniform distribution, where *I_A_* is the average current. We also modulate *I_A_* to keep the inhibitory cells at the threshold of firing even as 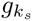 changes. At maximal 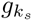 = 1.5 mS/cm^2^, *I_A_* is 1.0 *µ*A*/* cm^2^, and at minimal 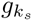 = 0 mS/cm^2^, *I_A_* is −0.14 *µ*A*/*cm^2^.

### Measures

To characterize network synchrony we employ a quantification dubbed the “Synchrony Measure” in previous work [18, 29, 30], a modified version of the measure developed in [31, 32]. This measures the degree of spike coincidence in a given neuron population for a specified time window. We define voltages for each neuron, *V_i_*(*t*), by convolving a Gaussian function with a series containing each neuron’s spike times over the window of interest. We obtain the population-averaged voltage 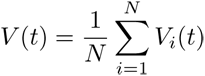, where N is the number of cells. Using *V_i_*(*t*) and *V* (*t*), we calculate the variance of the population voltage *σ* and variance of the individual neurons’ voltages *σ_i_* as

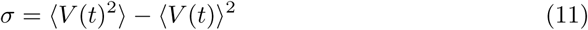

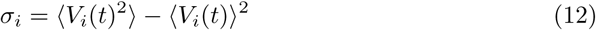

where ⟨⟩ denotes averaging over the time window. The Synchrony Measure S is then defined as

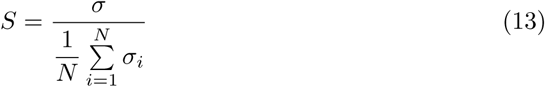

S assumes values from 0 to 1, with S = 0 denoting a completely asynchronous population and S = 1 denoting a completely synchronized population. We use rolling time windows without overlap to measure synchrony around different 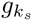 ranges, with the first window starting 1000 ms into the simulation when 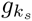 decline activates. For the simulations shown in Fig5, 6, and 7, 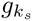 decline is set to 0.67 and the window lengths are set to 150 ms. For the simulations shown in Fig 4, the window lengths are set to either 50, 100, 200, or 400 ms, based on the four rates of 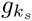 decline—networks with faster decline use smaller window lengths.

### Simulations

Code for all simulations is written in Python. We initialize each simulation with random initial conditions for each cell in the E-I network. *V* is selected from the uniform distribution (−62 mV, 22 mV), *n* and *h* from (0.2, 0.8), and *z* from (0.15, 0.25). Model equations are evaluated numerically using the fourth-order Runge-Kutta technique. We include synaptic current only after the first 100 ms of the simulation to allow initial transients to decay.

All raster plots are organized such that cells with higher *I_app_* have a lower neuron index and are placed closer to the bottom of the plots and vice versa. This allows for clearer visualization of the factors impacting temporal organization within individual bursts. For all figures with measures, we plot the average of the measures across ten independent simulations. We only compute measures based on network activity after the first 1000 ms, again to allow dynamics to stabilize.

### Data availability statement

There are no primary data in the paper; all materials are available at https://github.com/RichLabUConn/DynamicACh and the code is archived on Zenodo (DOI: 10.5281/zenodo.15641814).

## Results

### Network topology shapes response to dynamic cholinergic modulation

The cellular effects of varying levels of 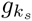 are illustrated in Fig 1; the extreme values of 1.5 and 0 mS/cm^2^ reflecting “low” and “high” levels of cholinergic modulation have been the primary focus in previous studies [29].

**Fig 1.**
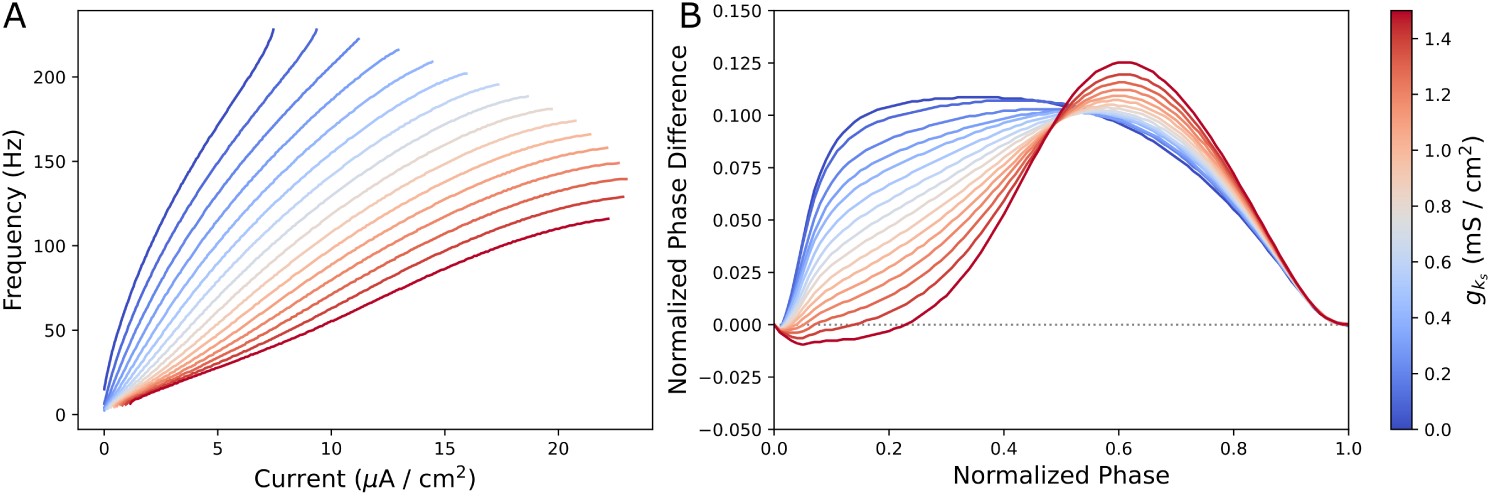
Changing intrinsic neuronal properties as a function of varying 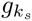. **A-B**: Frequency-Current (F-I) curves (Panel **A**) and phase response curves (PRCs, Panel **B**) for neurons with differing 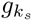. At 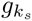 = 1.5 mS/cm^2^, the m-channel is maximally active and at 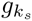 = 0 mS/cm^2^, the m-channel is fully blocked by cholinergic modulation. Panel **A** shows that neurons with lower 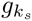 have steeper F-I curves, exhibiting higher excitability, and are capable of firing with lower current input. Panel **B** shows that neurons with higher 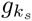 have a negative, phase delay region in response to an excitatory stimulus early in the neuron’s firing cycle. Cholinergic modulation induces a continuous shift in F-I curve and PRC properties.

Closure of the m-channel suppresses neuronal excitability (Fig 1A) and reduces the width of the phase delay region of the PRC (the duration of the window immediately following a spike in which an excitatory stimulus will delay the subsequent action potential) (Fig 1B). This phase delay region is responsible for a higher propensity for synchronization in populations of intra-connected excitatory neurons [33].

We first implemented dynamic cholinergic modulation, as realized in 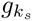 values varying linearly over time (detailed in the Materials and Methods), on E-I networks of two exemplar topologies: one with dominant inter-connectivity and the other with dominant intra-connectivity. These topologies were previously studied with tonic cholinergic modulation in [29].

As 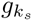 linearly decreases in these exemplar simulations (Fig 2), we note that the excitatory cells are driven to fire at increasingly higher frequencies until they all cease to fire (Fig 2A and C). Examining the voltage traces of the neurons confirmed that the cessation of firing was due to depolarization block. In Fig 2A, all inhibitory neurons also cease to fire, as a consequence of the heightened E-I connectivity in this network topology and thereby increased current supply to the inhibitory neurons.

**Fig 2.**
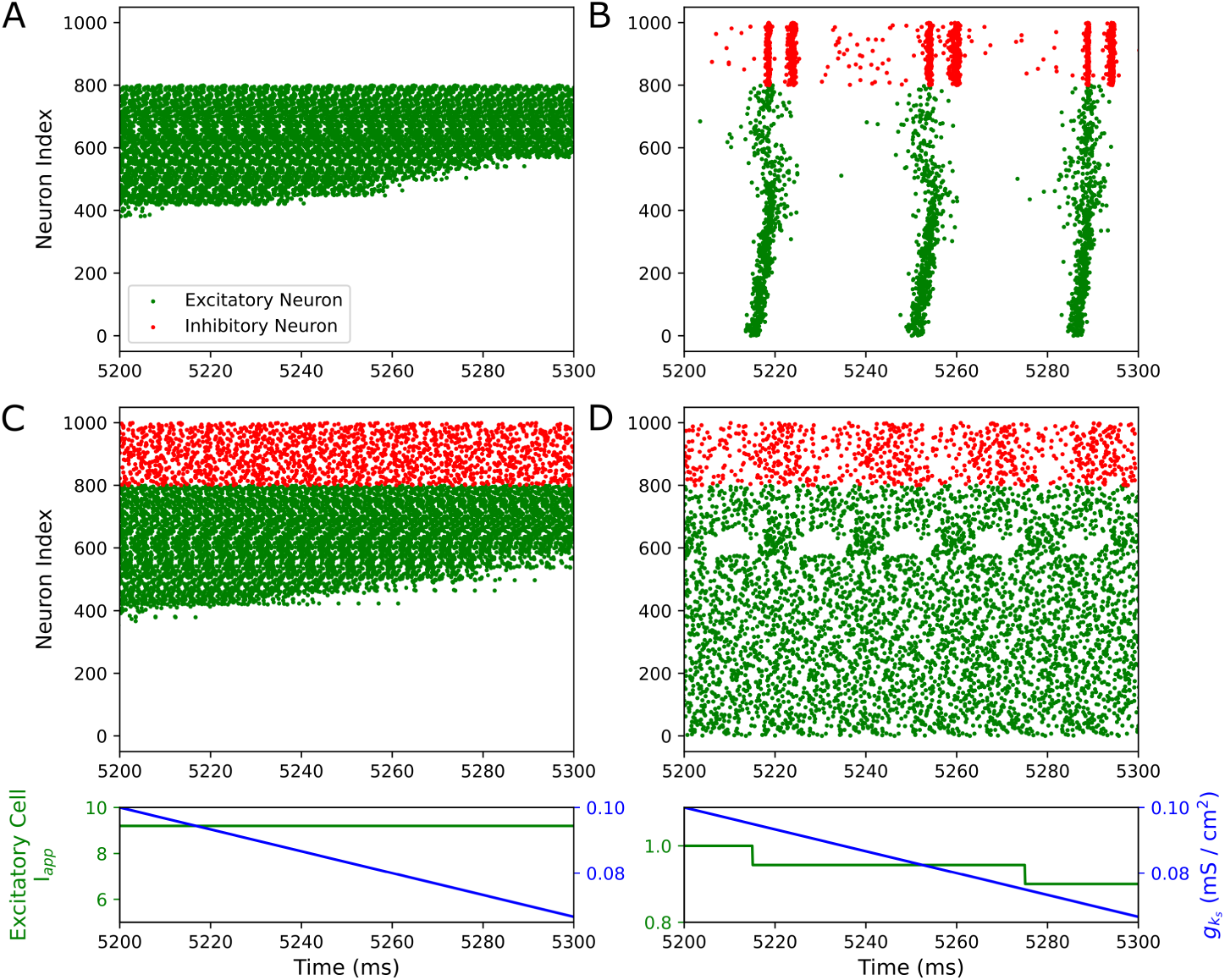
Dynamic m-channel blockade in E-I networks leads to depolarization block in the absence of concurrent external current modulation. **A-D**: Raster plots showing 100 ms periods of activity in E-I networks. Panels **A** and **B** are dominant inter-connectivity networks (synaptic weight parameters in mS/cm^2^: E-I, I-E = 0.00175, E-E = 0.0000625, I-I = 0.00025) and Panels **C** and **D** are dominant intra-connectivity networks (synaptic weight parameters in mS/cm^2^: E-I, I-E = 0.00025, E-E = 0.000125, I-I = 0.0005). Linear 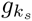 downregulation is carried out in all networks at rate 0.33 mS/cm^2^/s. 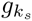 level is displayed below the raster plots, along with the average external current to an excitatory neuron, which is constant in Panels **A** and **C** and changing in Panels **B** and **D** to keep the firing frequency of an isolated average neuron approximately constant. Depolarization block occurs in response to declining 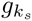 by default, with all excitatory neurons ceasing to fire (Panels **A** and **C**). However, when we account for higher neuron excitability induced by 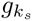 decrease by accordingly downregulating the external current, depolarization block does not occur (Panels **C** and **D**), preventing the biophysically unlikely phenomenon of cholinergic modulation inducing full depolarization block.

Termination of activity across all cells in a cortical microcircuit is an unrealistic response to cholinergic modulation from a physiological standpoint. These results arise disproportionately from the changes to the F-I curve as 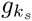 is scaled (Fig 1A), which impart a strong increase in cellular excitability that drowns out other effects of cholinergic modulation. To characterize less extreme effects of cholinergic modulation obscured by depolarization block in this first experiment, we introduced simultaneous modulation of the neurons’ current input (detailed in the Materials and Methods). This strategy controls for large changes in the baseline firing rate of isolated neurons while preserving the effects of cholinergic modulation on the gain of the F-I relationship and the changes to the PRC. Indeed, these modifications successfully prevent depolarization block in the example microcircuits (Fig 2B and D). Observing the stable end behaviors reached by both networks after this modification was implemented, i.e. after 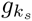 reached 0 mS/cm^2^, we found that only the dominant inter-connectivity network retained organized firing and that the dominant intra-connectivity network transitioned into asynchronous firing.

The end behavior seen here corresponds with findings regarding networks with tonic ACh [29] for both network topologies: networks with dominant intra-connectivity fire asynchronously at low ACh concentrations as they depend upon the phase delay region of the excitatory cells’ PRCs for synchronization, while networks with dominant inter-connectivity networks are able to synchronize independent of ACh. These results are affirmative support that a network whose 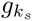 decreases dynamically will match the dynamics expected from a network with tonically low 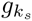 if the system is allowed to equilibrate. For the remainder of this manuscript, we will continue to modulate *I_ext_* in the manner discussed here.

We next asked whether the firing patterns exhibited in networks with dynamically changing 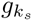 but *without* time to equilibrate would match the equilibrium dynamics of networks with an analogous constant 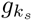 value. To do so, we compared the stable end behavior reached by networks simulated with 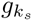 1.0 mS/cm^2^ (Fig 3A, C, and E) with the transient behavior of networks with time-varying 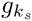 at a time window centered around 1.0 mS/cm^2^ (Fig 3B, D, and F).

**Fig 3.**
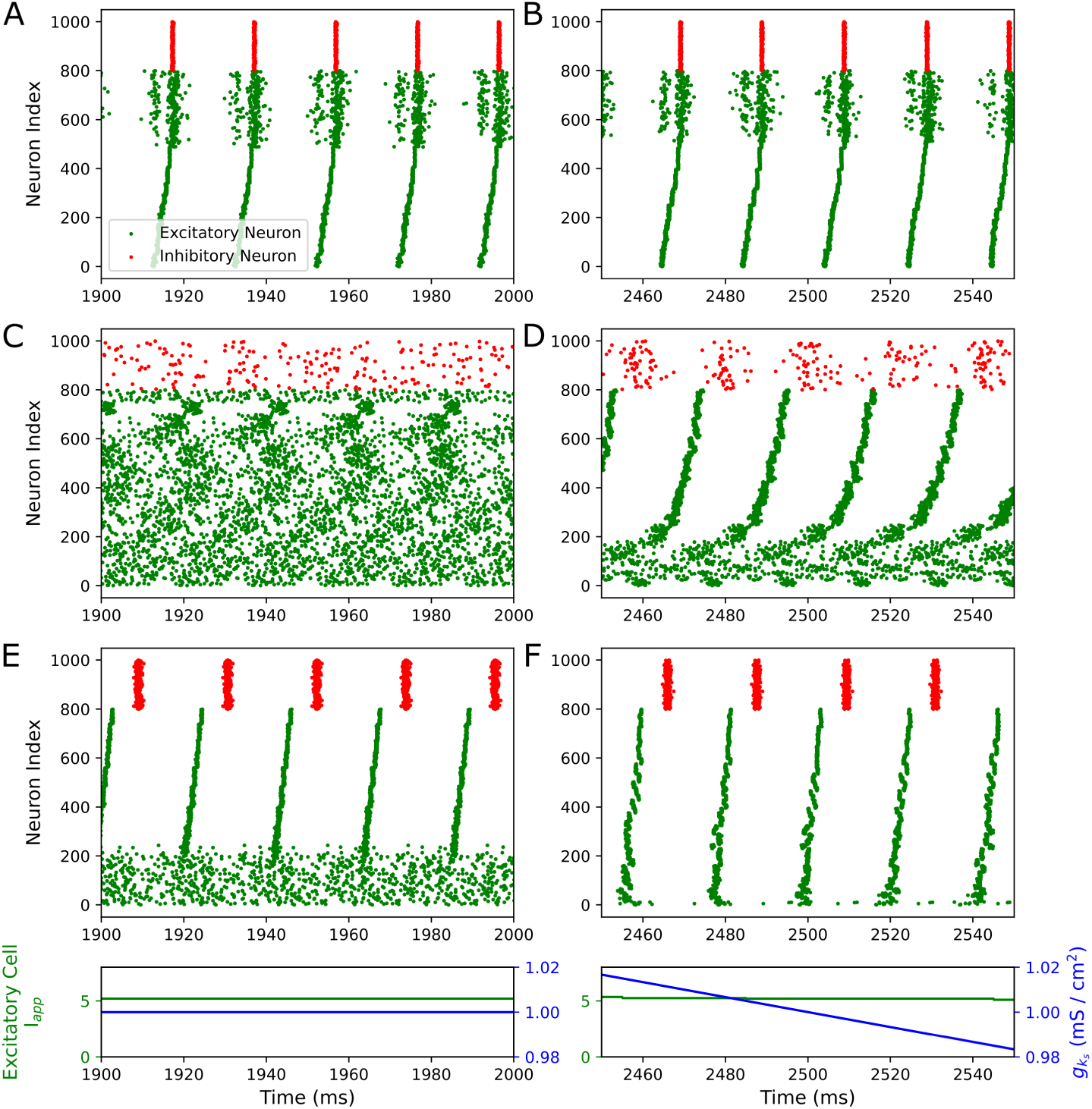
Differences in network activity caused by tonic versus dynamic cholinergic modulation arise preferentially in networks with dominant intra-connectivity. **A-F**: Raster plots showing 100 ms periods of activity in E-I networks, with constant 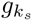 = 1.0 mS/cm^2^ in Panels **A**, **C**, and **E** and linearly declining 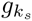 levels at rate 0.33 mS/cm^2^/s in Panels **B**, **D**, and **F**, at a window centered around 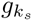 = 1.0 mS/cm^2^. Panels **A-B** have dominant inter-connectivity, and Panels **C-F** have dominant intra-connectivity, with parameters for synaptic weights as specified in Fig 2. Panels **E-F** have 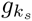 permanently set to 0 mS/cm^2^ for inhibitory cells. While dominant inter-connectivity networks show consistent organization of excitatory and inhibitory cells with dynamic (Panel **B**) or tonic (Panel**A**) 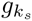 levels, dominant intra-connectivity networks display divergent patterning of the cells, as seen in the more synchronized behavior in Panels **D** and **F** compared to Panels **C** and **E**. The dominant intra-connectivity networks’ ability to fire synchronously is dependent upon cellular properties, which are influenced by the m-channel, explaining their tendency for altered firing patterns induced by changing 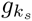.

In the dominant inter-connectivity networks, tonic (Fig 3A) versus dynamic cholinergic modulation (Fig 3B) induces no major differences in the firing patterns. In contrast, drastic differences arise in the dynamics between dominant intra-connectivity networks with tonic and phasic cholinergic modulation. The excitatory neurons in the examples with dynamic cholinergic modulation illustrated in Fig 3D and F (the former models joint cholinergic modulation of the excitatory and inhibitory neurons, the latter cholinergic modulation only in excitatory neurons) form distinctly more organized bursts in comparison to the corresponding examples in Fig 3C and E in which the 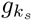 value is constant. Although we only showcase results from 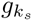 1.0 mS/cm^2^, similar relationships between tonic and dynamic models were found with other intermediate 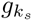 values in both network topologies during our investigations.

This preliminary investigation suggests that unique responses to time-varying ACh levels arise primarily in networks with dominant intra-connectivity. In networks with dominant inter-connectivity, consistent synchrony is independent of the intrinsic properties of individual neurons as a likely consequence of PING-like gating [29], intuitively explaining the similar dynamics arising from both dynamic and tonic changes to 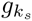. Meanwhile, the divergent behavior with dominant intra-connectivity networks is expected since these networks’ ability to synchronize is dependent upon cellular properties at high 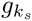 [29], motivating a more rigorous analysis of their behavior.

### Desynchronization in dominant intra-connectivity networks is sensitive to the rate of cholinergic modulation

To verify our intuition that only dominant intra-connectivity networks respond differentially to dynamic cholinergic modulation, we conducted simulations with varying rates of 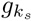 downregulation. Employing the Synchrony Measure (see Materials and Methods), we compared firing patterns between tonic and dynamic models across the entire 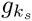 range (Fig 4). For the dynamically modulated networks, we compute the Synchrony Measure across moving time windows, allowing us to visualize the temporal evolution of the synchrony measure as 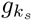 declines.

**Fig 4.**
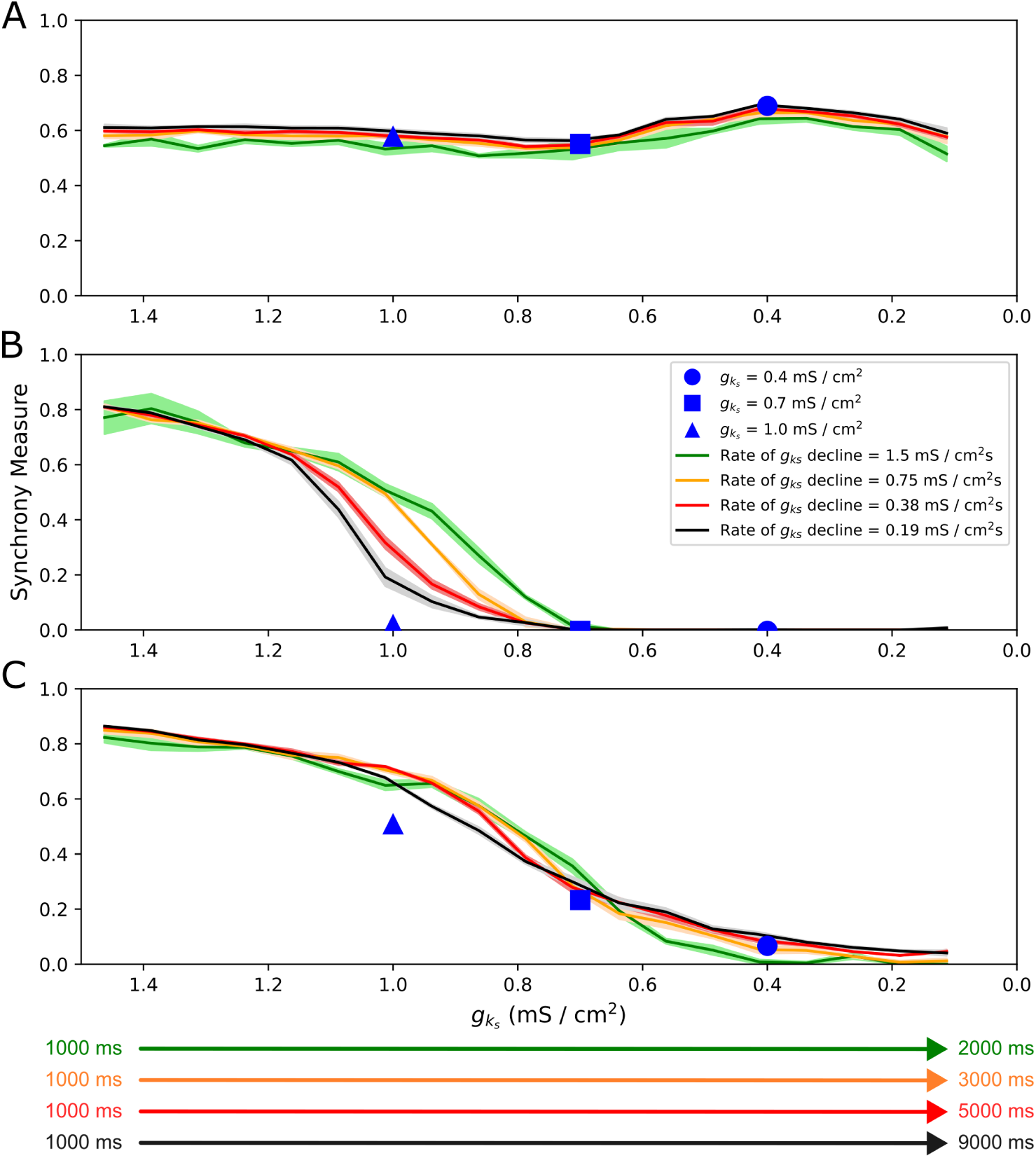
Rate of m-channel blockade affects the relationship between desynchronization and 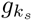 only in dominant intra-connectivity networks. **A-C:** Synchrony measure of excitatory cells plotted over 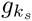 for networks with varying rates of 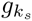 modulation and for networks with three illustrative tonic 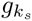 values. Panel **A** includes networks with dominant inter-connectivity and Panels **B-C** include networks with dominant intra-connectivity, with parameters for synaptic weights as specified in Fig 2. Panel **C** networks have 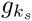 permanently set to 0 mS/cm^2^ for inhibitory cells. Color-coded arrows show the timecourse of 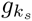 modulation from 1.5 to 0 mS/cm^2^ for each network. All measures plotted from tonic and dynamic networks are the mean based on 10 independent simulations, with *±* standard deviation shading displayed for dynamic networks. In Panel **A**, no substantial changes in synchrony are recorded in response to time-varying 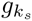. Conversely, in Panels **B-C**, desynchronization occurs, with slower rates of 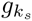 decline resulting in desynchronization starting at an earlier 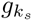 value. These observations indicate that strong inter-connectivity allows for synchronous network activity that is robust to cholinergic modulation regardless of its timecourse; however, for networks with dominant intra-connectivity, cholinergic modulation and its timescale affect the propensity for synchronous network activity over time.

Validating our predictions, the dynamics of dominant inter-connectivity networks is largely indifferent to the rate of modulation (Fig 4A). On the other hand, the dominant intra-connectivity networks are sensitive to the timecourse of cholinergic modulation. For 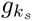 values between approximately 1.5 and 1.2 mS/cm^2^, the excitatory cells’ ability to retain organization is independent of the rate of modulation. As 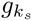 decreases past this point, the system moves towards asynchronous activity, with the behavior at a given 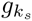 value sensitive to the rate of modulation. At a given value of 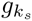 the dynamically modulated networks are at least as synchronized as analogous networks with a constant 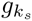, with the magnitude of this difference diminishing as 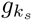 nears 0 mS/cm^2^. These patterns are broadly present in both Fig 4B and C, although in C, synchronous firing lasts longer for all the networks. This follows from the inhibitory cells being more excitable and more capable of forming tighter synchronized patterns with the absence of the m-current, which improves the efficacy of the inhibitory signaling to the excitatory population.

The comparisons between the dynamic and tonic networks as well as the similar dynamics exhibited at high 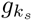 for all rates of modulation demonstrate that initially established firing patterns (i.e., at 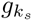 = 1.5 mS/cm^2^) play a potentially outsized role in the network dynamics during dynamic cholinergic modulation. The precipitous transition from synchronous to asynchronous dynamics as 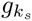 declines suggests that there may be a ‘critical’ concentration of ACh that is necessary to initiate desynchronization and that network behavior matches predictions from tonically modulated networks past this point. These analyses confirm that only networks with dominant intra-connectivity display behavior with our dynamic model of cholinergic modulation not predicted from analogous simulations with tonic cholinergic modulation. Their synchronization properties cannot be encapsulated as a function of the ACh concentrations at a given time alone, but rather depend upon the temporal evolution in ACh concentration—and in particular how that temporal evolution influences the history of firing activity. In the following, we continue study of networks with strong intra-connectivity using an exemplar 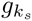 downregulation rate of 0.67 mS/cm^2^, which we found to be slow enough to allow for gradual changes in firing patterns while still remaining computationally tractable.

### The mechanism of ACh-driven desynchronization is especially sensitive to changes in E-I connectivity

With our focus on networks with strong intra-connectivity appropriately motivated, we further delineated the potentially interacting effects of dynamic cholinergic modulation and network topology on oscillatory activity. We independently varied E-I and I-E synaptic strengths to more precisely establish their effects on the dynamics driven primarily by intra-connectivity as described in detail above. For both inter-connectivity synapses we quantify network dynamics as we gradually increase their weight, up to and including the value used previously in our “dominant inter-connectivity” network. We conduct this experiment in networks with and without cholinergic modulation of the inhibitory neurons, accounting for potential variations in m-channel expression among inhibitory interneurons in different brain regions [34] (detailed in Discussion).

Variations in E-I connectivity have acute and non-monotonic implications for the duration of synchronous activity in the network. This is illustrated in Fig 5A: increases to the E-I synaptic weight up to 0.0008125 mS/cm^2^ delay desynchronization in the network, whereas increases above this threshold instead hasten desynchronization, causing it to occur at higher 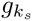 values. By inspecting raster plots we found that desynchronization in networks with an E-I synaptic weight below this threshold (which includes the default dominant intra-connectivity network studied in previous sections) starts with faster-firing cells (Fig 5C1). Meanwhile, above this threshold, we observe a distinct pattern of desynchronization that starts with slower-firing cells (Fig 5C2). We hypothesized that these dichotomous patterns were indicative of two distinct mechanisms of ACh-modulated desynchronization, demarcating two regimes (red and blue boundaries in Fig 5A) determined by the strength of E-I connectivity.

**Fig 5.**
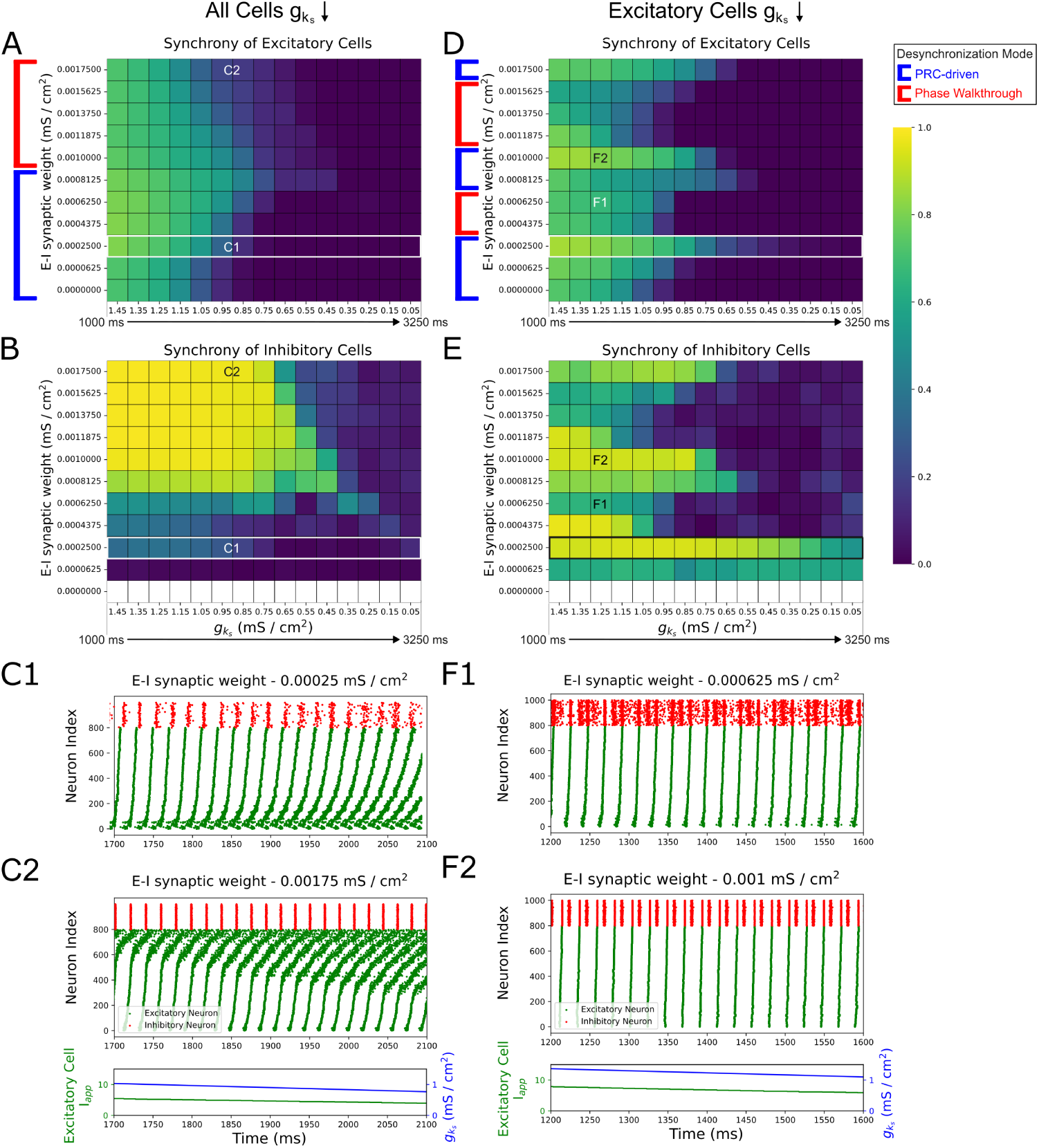
E-I connectivity controls speed and mechanisms of ACh-induced desynchronization in dominant intra-connectivity networks. **A, B, D, E**: Heatmaps of excitatory (Panels **A** and **D**) and inhibitory (Panels **B** and **E**) synchrony averaged over 10 independent simulations in networks of varying E-I connectivity. Cholinergic modulation is implemented with 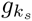 decline rate 0.67 mS/cm^2^/s, carried out for all cells (Panels **A-B**) or for only excitatory cells while inhibitory cells’ 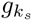 is permanently set to 0 mS/cm^2^/s (Panels **D-E**). E-E, I-I, and I-E synaptic weights are fixed at 0.000125 mS/cm^2^, 0.0005 mS/cm^2^, and 0.00025 mS/cm^2^ respectively, with the default dominant intra-connectivity networks denoted with special white / black borders. **C, F**: Raster plots showing 400 ms periods of activity selected from networks of differing synaptic weights as labeled, for networks with modulation for all cells (Panel **C**) and for only excitatory cells (Panel **F**). Variations in E-I connectivity induce non-monotonic shifts in the duration of synchronous firing that are underscored by changes in the mode of desynchronization (Panels **A** and **D**). Panel **C** showcases how these distinct modes of desynchronization manifest: PRC-driven desynchronization occurs due to the attenuation of the phase delay region in the excitatory cells’ PRC (Panel **C1**), while phase walkthrough-driven desynchronization which occurs due to the inhibitory bursts arriving progressively earlier and disrupting the firing times of the slower excitatory cells (Panel **C2**). Panel **F** demonstrates the development of multiple inhibitory population bursts per excitatory population burst, a factor that contributes to switches in desynchronization mode seen in Panel **D**.

The pattern of desynchronization below the threshold E-I synaptic weight is, as for our dominant intra-connectivity network, viably explained by the changes to the excitatory cells’ PRC properties induced by 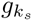 decline and is thus termed “PRC-driven”. Although all the networks in this range desynchronize in this manner, the increase in E-I connectivity delays desynchronization by better entraining the inhibitory population. This is evidenced in the qualitative relationship between the magnitude of the initial synchrony in the inhibitory population (leftmost column of Fig 5B) and the duration of excitatory synchronous activity (blue-bounded networks in Fig 5A). This occurs as more synchronous inhibitory bursts create a stronger and more uniform inhibitory signal to each of the excitatory cells, minimizing the cell-to-cell variability in this signal and strengthening any resulting gating effects.

As a result, in spite of weak I-E synapses, desynchronization is delayed through a mechanism with some similarity to the PING mechanism. Unlike with PING-driven networks however, the inhibitory signaling here only preserves existing synchronous activity rather than serving as the impetus for its formation. This is confirmed when we set the strength of the E-I synaptic weight to 0 mS/cm^2^: while inhibitory cells are inactive (white bottom row of Fig 5B), we find that the excitatory cells’ synchrony shows no apparent difference from the lowest non-zero E-I synaptic weight (network with E-I synaptic weight 0.0000625 mS/cm^2^ in Fig 5A).

We attribute the phenomenon in the second regime (red-bounded networks in Fig 5A), where desynchronization begins with slower firing cells, to “phase walkthrough” (borrowing terminology from previous PING literature [22]). Raster plot inspection revealed that once a sufficiently low 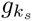 value is reached in these networks, high E-I connectivity leads inhibitory bursts to arrive progressively earlier relative to the excitatory bursts, breaking the phase-lock between the excitatory and inhibitory populations—the defining characteristic of phase walkthrough. When the inhibitory bursts arrive early enough that they precede and disrupt the spiking of the slower firing excitatory cells, signaling between the E and I populations becomes weaker with more cell-to-cell variability, initiating desynchronization of the excitatory population.

E-I connectivity scaling when cholinergic modulation is restricted to excitatory cells (Fig 5D-E) produces similar outcomes, albeit with multiple non-monotonic shifts in the duration of synchronous activity. Correspondingly, we note a tight set of “optimal” E-I synaptic weights (0.00025, 0.0008125, 0.001, 0.00175 mS/cm^2^ in Fig 5D) at which relatively long-lasting synchronous activity is present. We confirmed that these E-I synaptic weights lie at points of transition between PRC-driven desynchronization regimes and phase walkthrough-driven desynchronization regimes by inspecting the raster plots and identifying whether the desynchronization began with faster or slower-firing cells.

We explain this behavior through the higher excitability of the inhibitory cells locked at 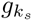 = 0 mS/cm^2^. This excitability allows each inhibitory cell to respond with multiple spikes in response to each excitatory burst as E-I connectivity is increased (Fig 5F); notably, this phenomenon was never observed in the scenario described in Fig 5A-C. However, the number and timing of responses from each inhibitory cell does not change in a uniform manner: for example, at 0.000625 mS/cm^2^ (Fig 5F2), the inhibitory clusters that follow each excitatory burst are highly temporally disorganized, but at 0.001 mS/cm^2^, we observe two well-defined, tight bursts in response to each excitatory burst. This general trend is evidenced in the leftmost column of Fig 5E, where we observe that the initial synchrony of the inhibitory population changes in a non-uniform manner with increased E-I synaptic weight.

We propose that the aforementioned optimal E-I synaptic weights occur when the inhibitory cells are able to organize into well-defined, tight bursts and are thereby able to delay desynchronization through effective gating. For example, at 0.00025 mS/cm^2^, the initial inhibitory synchrony is high (leftmost column of Fig 5E), and the excitatory cells are able to stay synchronized for a relatively longer duration (Fig 5D). Further increase in the E-I synaptic weight to 0.0004375 mS/cm^2^ results in earlier desynchronization through phase walkthrough, despite high initial inhibitory synchrony, as was also the case with the phase walkthrough regime in Fig 5A and Fig 5B. As E-I connectivity is scaled past an E-I synaptic weight at which phase walkthrough occurs, the increase in the inhibitory cells’ firing frequency, coupled with the strong I-I connectivity slowing the inhibitory bursts, appears to allow a return to the regime where network desynchronization is driven by PRC effects and phase walkthrough does not occur (for example, 0.0008125 mS/cm^2^ case in Fig 5D and E).

Thus, E-I connectivity plays an important role in controlling the manner and rate of desynchronization by controlling the timing and strength of the inhibitory signaling. Subtle differences in this connectivity may drastically alter how a network responds to cholinergic modulation, showing that network topology works together with ACh-induced changes in cellular properties to direct dynamic network behavior.

In contrast, strengthening I-E connectivity uniformly delays PRC-driven desynchronization (Fig 6A and Fig 6D), with no cases of phase walkthrough, as verified by inspecting the raster plots and finding that desynchronization consistently began with the faster firing cells. The delay in PRC-driven desynchronization here may be attributed to the increased I-E synaptic weights strengthening inhibitory signaling to the excitatory cells and consequently strengthening gating effects; as was noted above for the PRC-driven desynchronization regime of Fig5A, this gating has some similarities to the PING mechanism.

**Fig 6.**
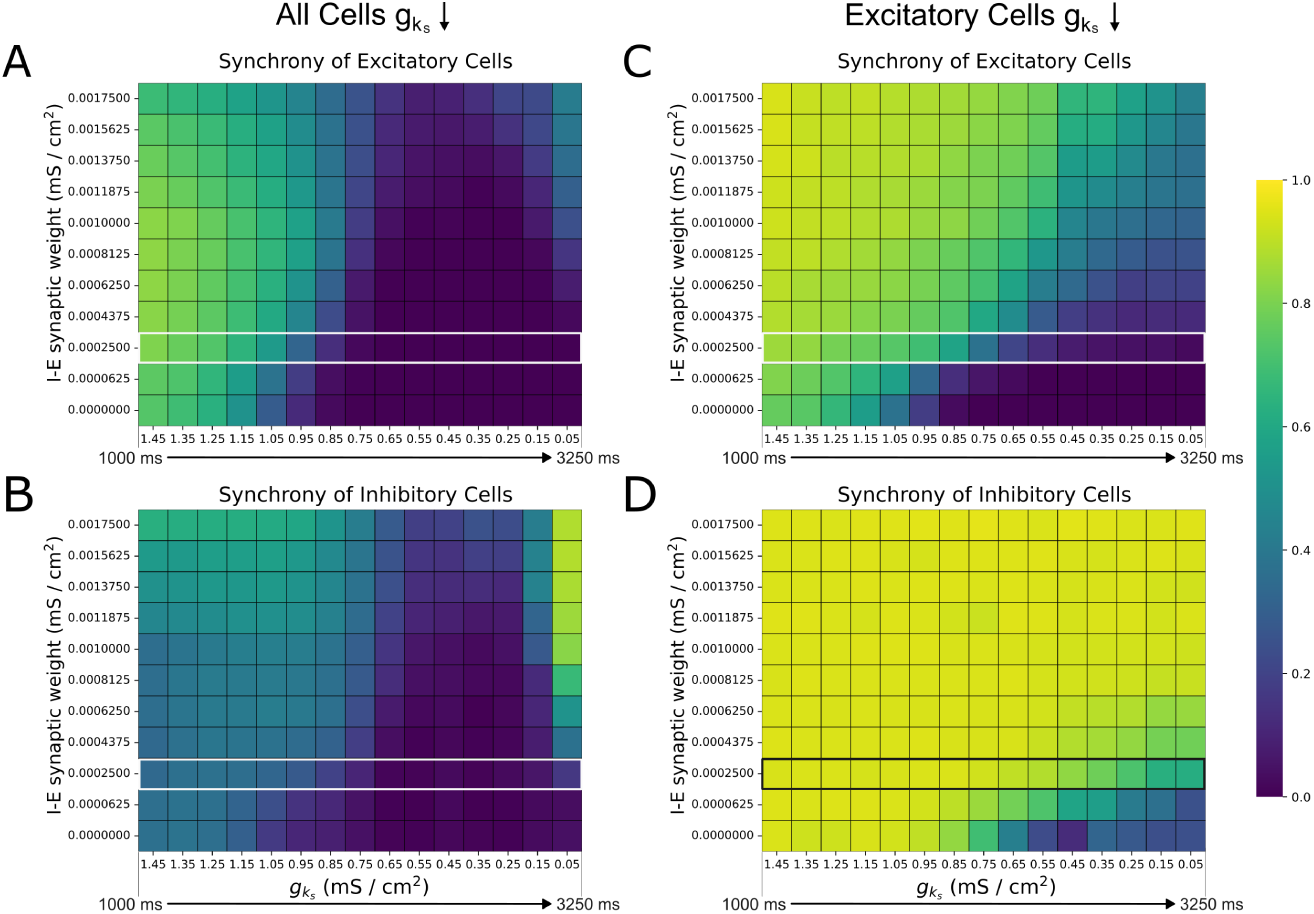
I-E connectivity prolongs synchronous firing for dominant intra-connectivity networks subjected to cholinergic modulation.. **A, B, C, D**: Heatmaps of excitatory (Panels **A** and **C**) and inhibitory (Panels **B** and **D**) synchrony averaged over 10 independent simulations in networks of varying I-E connectivity. Cholinergic modulation is implemented with 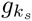 decline rate 0.67 mS/cm^2^/s, carried out for all cells (Panels **A-B**) or for only excitatory cells while inhibitory cells’ 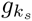 is permanently set to 0 mS/cm^2^/s (Panels **C-D**). E-E, I-I, and I-E synaptic weights are fixed at 0.000125 mS/cm^2^, 0.0005 mS/cm^2^, and 0.00025 mS/cm^2^ respectively, with the default dominant intra-connectivity networks denoted with special white / black borders. Strengthening I-E connectivity gradually increases network propensity for synchronization (Panels **A** and **D**) and enables reemergence of synchrony at low 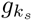 (Panel A).

Notably, in Fig 6A, weakly synchronous activity (Synchrony Measure values of approximately 0.4 at I-E synaptic weight 0.001375 mS/cm^2^ and above) reemerges after desynchronization for sufficiently high I-E connectivity. This resurgence occurs at low 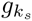 values, where the inhibitory cells are more excitable and better able to reorganize into synchronous bursts (Fig 6B), allowing them to re-initiate oscillatory network activity. Conversely, when cholinergic modulation is restricted to the excitatory cells and inhibitory cells’ 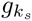is locked at 0 mS/cm^2^ (Fig 6C and D), highly synchronized inhibitory bursts are consistent through time for most I-E synaptic weights, preventing the excitatory cells from desynchronizing at moderate values of 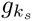.

Thus, while both E-I and I-E synapses mediate ACh-driven desynchronization, changes in E-I connectivity exert stronger control over the desynchronization rate while changes in the I-E connectivity have a weaker and more uniform effect. Previous research [29] suggests that the PING-driven effects of strong inter-connectivity would dominate the ACh-driven effects of strong intra-connectivity when all synaptic weights are strong. We validate this in Fig 7: in networks with simultaneously strong intra- and inter-connectivity (purple networks in Fig 7A and B in which each synaptic weight matches the value in the associated “dominant” connectivity network), the stability created by strong inter-connectivity promotes activity qualitatively matching that of dominant inter-connectivity networks (blue networks in Fig 7A and B). Indeed, despite some quantitative differences and fluctuations in the synchrony measure, these networks retain oscillatory firing for all 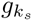 values.

**Fig 7.**
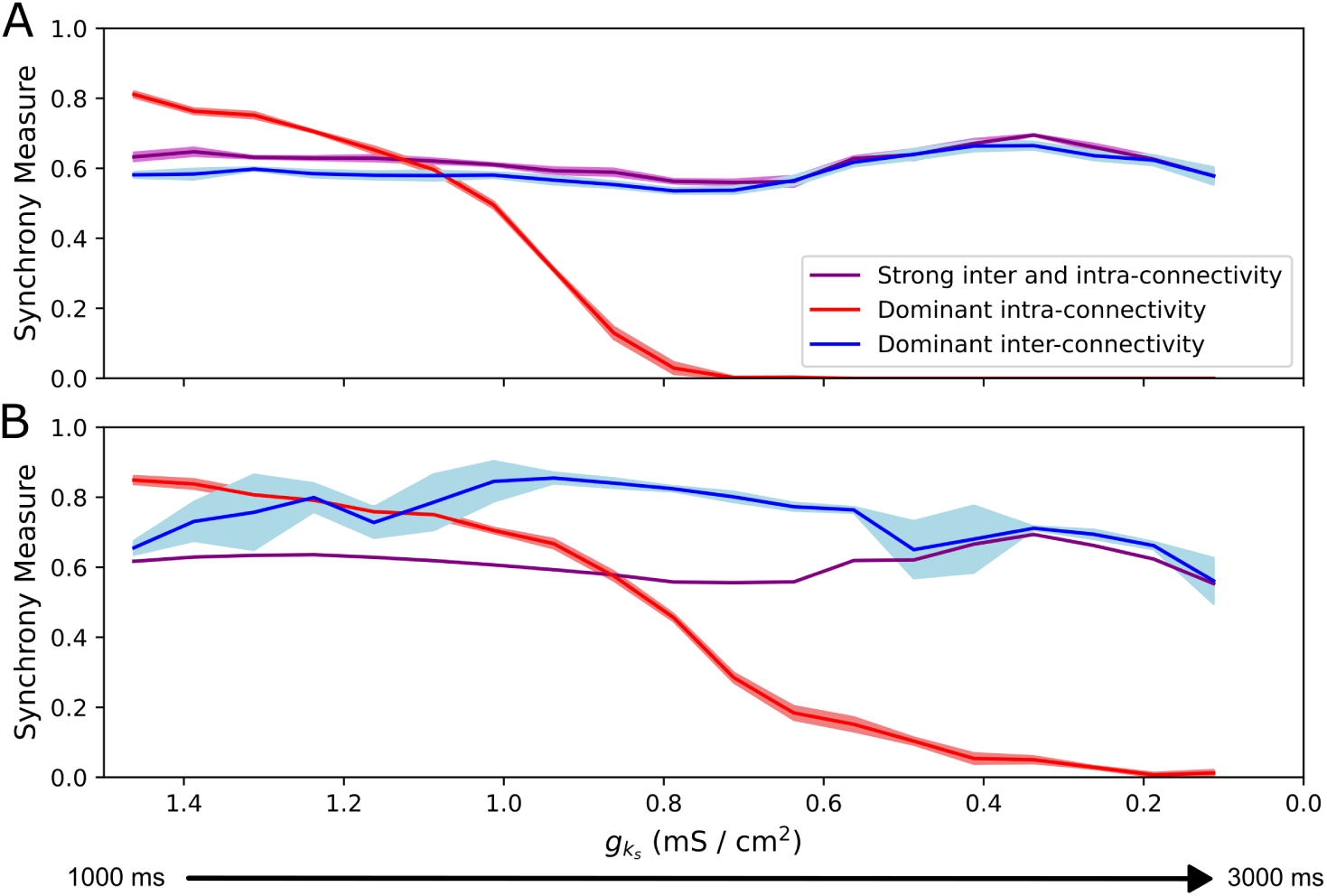
In systems with both strong inter- and intra-connectivity, network response is dominated by interconnectivity-driven effects. **A-B**: Synchrony measure of excitatory cells plotted over 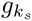 for networks with varying network topologies and cholinergic modulation for all cells (Panel **A**) or for only excitatory cells (Panel **B**). Synaptic weight parameters for dominant intra-conectivity and dominant inter-connectivity synaptic weights are as specified in Fig 2 and as follows for strong inter and intra-connectivity networks: E-I, I-E = 0.00175, E-E = 0.000125, I-I = 0.0005 mS/cm^2^. All measures plotted from tonic and dynamic networks are the mean based on 10 independent simulations, with *±* standard deviation shading displayed. When E-I and I-E synaptic weights are both high, network behavior is largely uninfluenced by cholinergic modulation, irrespective of strong intra-connectivity.

## Discussion

Through study of an E-I network whose component neurons are subject to dynamic cholinergic modulation of the m-channel, we here delineate how time-varying ACh concentrations and network topology interact to influence firing patterns in *in silico* cortical microcircuits. Specifically, we determine that oscillations driven by strong connectivity within the excitatory population are disrupted by dynamic cholinergic modulation unlike oscillations driven by strong connectivity between the excitatory and inhibitory populations, with the former networks giving rise to unique dynamics sensitive to the timecourse of ACh evolution. Consequently, networks in which population intra-connectivity dominates over population inter-connectivity are most influenced by time-varying ACh levels, with small changes to the E-I connectivity in such networks having disproportionate effects on network synchrony. Collectively, these analyses highlight that the traditional view of cholinergic modulation as acting on long neuromodulatory timescales may obscure relevant dynamics arising when ACh levels vary in more temporally precise fashions.

This computational work is motivated by emerging experimental evidence that ACh acts on millisecond-level timescales via muscarinic receptors in the brain [6, 11], a phenomenon that is understudied in contemporary computational literature. The primary source of cholinergic projections to the cortex, the basal forebrain, is known to deliver precisely timed phasic cholinergic transients [7, 35]. This signaling has been found to alter cortical firing patterns, promoting a transition from low-frequency rhythmic activity to desynchronized firing as seen in local field potentials and EEG recordings [4, 36]. Such ACh-induced desynchronization arises not only during sleep-wake cycles, but also rapidly, on the timescale of seconds or shorter, during tasks that engage sensory and motor skills [37, 38]. The work presented here represents a pivotal first step towards delineating *in silico* mechanisms by which such experimentally-observed effects arise.

Based on the analyses presented here and previous computational explorations [29], we hypothesize that ACh facilitates rapid desynchronization in networks whose oscillations manifest through intrinsic cellular properties dependent upon m-current activity [33], but not in networks with oscillations arising from PING-like gating mechanisms [22, 39]. This desynchronization occurs through two distinct mechanisms: one dependent upon changes to the excitatory cells’ PRCs [12] and the other through phase walkthrough effects attributed to excessive excitation of inhibitory cells [22].

While our study focuses on desynchronization of cortical oscillations triggered by increases in ACh, in some experimental settings cholinergic transients have a synchronizing effect—for instance, in the formation of gamma rhythms in the prefrontal cortex during attentional performance tasks such as cue-detection [40]. This phenomenon has been captured computationally in E-I networks with pulsatile time-varying modulation of the m-current [28]. The distinct experimental phenomena captured by our model and the work of [28] is reflected in an important *in silico* nuance: [28] does not control for increases in neuronal baseline firing frequency induced by the effect of 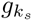 modulation on the F-I curve (as done in our work) given the impermanent nature of their modeled ACh increases. This implies that that such synchronization effects are likely more dependent on the heightened excitability induced by F-I curve modulation, as suggested in [28] itself, than the PRC modulation effects of primary focus in our study. This not only reconciles these seemingly disparate findings regarding cholinergic modulation’s influence on network synchrony, but highlights the need for *in silico* study to parse out the most salient effects of cholinergic modulation on network dynamics in varying settings.

We note that we examined networks with and without an active m-current in inhibitory cells, in consideration of the conflicting evidence regarding muscarinic receptor prominence among parvalbumin interneurons in the visual cortex [34]. We found that networks with cholinergic modulation for inhibitory cells may allow for partially synchronous firing in high ACh networks, a result predicted by previous study of the m-channel’s influence on patterning in purely inhibitory networks [18]. Interestingly, both desynchronization modes we defined arose in networks with exclusive cholinergic modulation of excitatory cells; in particular, phase walkthrough-driven desynchronization can occur even without modulating 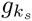 for the inhibitory cells. These findings indicate that the primary conclusion of this work—that the effects cholinergic modulation at millisecond time scales are strongly dependent upon network topology—is generalizable to brain regions with differing m-channel expression on inhibitory interneurons.

This study purposefully focuses on cholinergic modulation through muscarinic receptors, motivated in part by the extensive computational literature focused on the muscarinic receptor-gated m-channel [24–28] as well as experimental evidence that muscarinic receptors play a crucial role in desynchronizing cortical oscillations [11, 36] and in regulating cognitive functions such as attentional modulation [41]. Of particular relevance are findings from the study of vagal nerve stimulation (VNS) that some of this treatment’s therapeutic effects are driven by its activation of cholinergic pathways through the basal forebrain [42, 43]. Although VNS mechanisms of action are not yet fully understood [42–44], there is emerging evidence indicating muscarinic pathways may play a role in realizing its benefits. This is perhaps best described in pre-clinical work [45] finding that application of the muscarinic antagonist scopolamine greatly mitigates VNS’ ability to desynchronize noise-driven spikes in the auditory cortex. Rigorously deciphering how a time-varying, VNS-driven cholinergic signal affects cortical activity will be imperative not only to advance our general knowledge regarding VNS, but also to understand its efficacy in the newly FDA-approved use of VNS in post-stroke motor rehabilitation: this treatment requires precise temporal pairing of VNS during motor rehabilitation [46], echoing findings in rodent studies [47–50].

Our *in silico* work represents a vital proof of principle that features including the rate of change of ACh concentration and network topology affect how a system responds to a time-varying cholinergic signal, which could be particularly relevant to post-stroke VNS-paired motor rehabilitation given the experimental and clinical results described above. Such studies will likely require additional biophysical details to maximize clinical applicability, perhaps including spatial heterogeneity in cholinergic modulation (in recognition of evidence suggesting that sites of ACh transmission in the neocortex are highly localized [6, 8]) as well as an inclusion of ACh’s effect on nicotinic receptors. Nonetheless, the idealized computational results presented here represent strong motivation for these future studies delineating the effects of a wide-range of time-varying cholinergic signals, including signals derived from VNS.

